# Adolescent intermittent ethanol exposure induces sex-dependent divergent changes in ethanol drinking and motor activity in adulthood in C57BL/6J mice

**DOI:** 10.1101/2020.04.28.066472

**Authors:** Antoniette M. Maldonado-Devincci, Joseph G. Makdisi, Andrea M. Hill, Renee C. Waters, Nzia I. Hall, Mariah Shobande, Anjali Kumari

**Author notes:** To whom correspondence should be addressed: Antoniette M. Maldonado-Devincci, Ph.D. North Carolina A&T State University Department of Psychology, 1601 East Market Street, New Science Building Room 360 Greensboro, North Carolina 27411. Funded by the National Center for Advancing Translational Sciences, National Institutes of Health, through Grant KL2TR002490 and the North Carolina Agricultural and Technical State University Division of Research and Economic Development (AMD).

## Abstract

With alcohol readily accessible to adolescents, its consumption leads to many adverse effects, including impaired learning, attention, and behavior. Adolescents report higher rates of binge drinking compared to adults. Adolescents are also more prone to substance use disorder during adulthood due to physiological changes during the adolescent developmental period. We used C57BL/6J male and female mice to investigate the long-lasting impact of binge ethanol exposure during adolescence on voluntary ethanol intake and open field behavior during later adolescence and in young adulthood. The present set of experiments were divided into four stages: (1) chronic intermittent vapor inhalation exposure, (2) abstinence, (3) voluntary ethanol intake, and (4) open field behavioral testing. During adolescence, male and female mice were exposed to air or ethanol using an intermittent vapor inhalation with repeated binge pattern ethanol exposure from postnatal day (PND) 28–42. Following this, mice underwent abstinence during late adolescence from PND 43–49 (Experiment 1) or PND 43–69 (Experiment 2). Beginning on PND 49–76 (Experiment 1) or PND 70–97 (Experiment 2), mice were assessed for intermittent voluntary ethanol consumption using a two-bottle drinking procedure over 28 days. Male mice that were exposed to ethanol during adolescence showed increased ethanol consumption during later adolescence (Experiment 1) and in emerging adulthood (Experiment 2), while the female mice showed decreased ethanol consumption. These data demonstrate a sexually divergent shift in ethanol consumption following binge ethanol exposure during adolescence and differences in open field behavior. These data highlight sex-dependent vulnerability to developing substance use disorders in adulthood.

**Significance Statement:** Currently, it is vital to determine the sex-dependent impact of binge alcohol exposure during adolescence, given that until recently females have largely been ignored. Here we show that adolescent male mice that are exposed to binge ethanol during adolescence show long-term changes in behavior in adulthood. In contrast, female mice show a transient decrease in ethanol consumption in adulthood and decreased motor activity spent in the center zone of the open field test. Male mice appear to be more susceptible to the long-term changes in ethanol consumption following binge ethanol exposure during adolescence.

## 1. Introduction

The adolescent developmental period across mammalian species has been associated with various age-related physiological, hormonal, neuronal, cognitive, and behavioral changes during the transition from childhood to adulthood (for review see (Spear, 2000)). Behaviorally, adolescents show increases in social behavioral motivations, risk-taking, and novelty-seeking, which may contribute to the initiation and escalation of alcohol use during this time (Patrick et al., 2011; Spear, 2016). On average, adults drink more often than adolescents, however, both humans and rodent models have shown that per occasion adolescent alcohol consumption is much higher than that of adults (Doremus et al., 2005; SAMHSA, 2008).

Adolescent onset drinking before age 15 in humans has been associated with several fold higher drinking in adulthood compared to those that waited longer to initiate alcohol consumption (Dawson et al., 2008). Binge drinking during adolescence leads to higher rates of substance use disorder symptoms by the age of 35, including dependence on alcohol, abuse, disease, and death (Patrick et al., 2011). Although various factors contribute to alcohol abuse during adolescence, early and repeated heavy alcohol consumption is associated with an increased propensity for elevated intake of alcohol in later life in both humans and rodents (for review see (Spear, 2016)). Consequently, episodic heavy or binge drinking has proven to be a major point of concern in understanding adolescent alcohol drinking behaviors (Johnston et al., 2019).

Animal models not only mitigate the ethical implications of using humans but also control for genetic and environmental influences when studying the long-term effects of adolescent alcohol behaviors on adulthood alcohol consumption (Ehlers et al., 2013). In previous rodent studies, the impact of adolescent intermittent ethanol (AIE) exposure on adult drinking patterns and behaviors has varied. AIE has regularly been shown to increase alcohol drinking, social anxiety, and loss of behavioral flexibility in adulthood in rats (Crews et al., 2019; Gilpin et al., 2012; Hargreaves et al., 2009; A.M. Maldonado-Devincci et al., 2010; Antoniette M. Maldonado-Devincci et al., 2010; Pascual et al., 2009), while others found no change in alcohol consumption following adolescent ethanol exposure (Nentwig et al., 2019; Slawecki and Betancourt, 2002). Other studies in rats reported long-term negative effects on cognitive flexibility, including impairments in spatial learning and completing aversive tasks (Acheson et al., 2013; Miller et al., 2017). These differences in previous work are likely due to varied experimental exposure patterns, caging methods, and /or administration procedures of alcohol.

Several models of AIE exposure have been used in both rats and mice, including voluntary consumption in genetically bred mouse and rat strains, forced gavage exposure, intermittent intraperitoneal injections, forced ethanol consumption with only one bottle of ethanol presented, drinking in the dark, etc. (Acheson et al., 2013; Crews et al., 2019; Gilpin et al., 2012; A.M. Maldonado-Devincci et al., 2010; Miller et al., 2017; Slawecki and Betancourt, 2002; Thiele and Navarro, 2014).

Another method that has recently been used for AIE exposure is intermittent vapor ethanol exposure (Barker et al., 2017; Gass et al., 2014; Nentwig et al., 2019; Trantham-Davidson et al., 2017). This method exposes rats to two days of intermittent vapor inhalation to maintain high and elevated blood ethanol concentrations (BEC) (Gass et al., 2014). These studies in rats showed that AIE alters PFC-related decision making and behavioral control, but not ethanol-mediated behavior in adulthood (Centanni et al., 2017; Gass et al., 2014; Nentwig et al., 2019; Trantham-Davidson et al., 2017). Other studies using moderate to high chronic vapor ethanol exposure in male rats (e.g., BECs between 150–225 mg/dl) have shown hypoactivity in locomotion tests, increased anxiety-like behavior in the open field test, depressive-like behavior, and lower body weight following exposure (Ehlers et al., 2013; Slawecki et al., 2004; Walker et al., 2010). Other work in rats showed that AIE vapor exposure combined with intermittent ethanol consumption during adolescence increases drinking in adulthood (Amodeo et al., 2018).

The vapor inhalation model of ethanol dependence increases ethanol consumption in ethanol-dependent mice (Becker and Lopez, 2004; Carrara-Nascimento et al., 2013; Dhaher et al., 2008; Griffin et al., 2009; Jury et al., 2017; Lopez and Becker, 2005; Metten et al., 2010). The intermittent vapor inhalation AIE model used in adolescent mice showed that ethanol exposure induces a conditioned taste aversion in adulthood (Diaz-Granados and Graham, 2007). Other work has shown that chronic intermittent ethanol (CIE) vapor exposure increased ethanol drinking in both adolescent and adult male mice when ethanol consumption was measured between cycles (Carrara-Nascimento et al., 2013).

To date, most studies have exposed animals to alcohol/drugs during adolescence compared to adulthood, where animals are exposed at different ages and the withdrawal period is the same number of days (Galaj et al., 2020; Pascual et al., 2009; Tambour et al., 2008). In the present work, we exposed mice at the same age during adolescence and the length of withdrawal was varied to allow us to understand the impact of short-term and protracted withdrawal on subsequent ethanol consumption and open field behavior. This is similar to recent work that varied the withdrawal period to assess short-term and long-term changes in ethanol consumption and anxiety-like behavior (Leonardo Jimenez Chavez et al., 2020; Salguero et al., 2020). Recently it was shown that adolescent binge ethanol exposure decreased locomotor activity in the open field test in rats (Salguero et al., 2020) and induced long-term changes in anxiety-like behavior in adulthood in mice (Leonardo Jimenez Chavez et al., 2020).

The present experiment was designed to use a model of binge ethanol exposure using the vapor inhalation exposure paradigm during adolescence to examine sex differences in the long-term consequences in voluntary unsweetened ethanol intake in later adolescence and in adulthood in male and female mice. Specifically, we exposed adolescent (PND 28–42) C57BL/6J mice to adolescent intermittent ethanol (AIE) vapor exposure and measured voluntary ethanol intake (1) one week following the initial exposure while the mice were still adolescents and (2) four weeks following the AIE exposure when the mice were young adults (PND 70) and assessed subsequent locomotor activity. Similar to other work conducted in rats (Crews et al., 2019; Gilpin et al., 2012; Hargreaves et al., 2009; A.M. Maldonado-Devincci et al., 2010; Antoniette M. Maldonado-Devincci et al., 2010; Pascual et al., 2009), we expected adolescent intermittent ethanol exposure to increase voluntary ethanol intake in adulthood in both male and female mice. Additionally, we expected female mice to consume more ethanol compared to male mice (Amodeo et al., 2018; Jury et al., 2017; Morales et al., 2015; Tambour et al., 2008). Finally, we expected to see lower overall locomotor activity and greater thigmotaxis in mice exposed to ethanol during adolescence compared to control mice exposed to air.

## 2. Methods

### 2.1. Subjects

Adolescent male and female C57BL/6J mice (n=10–13/group) were obtained from Jackson Laboratories (Bar Harbor, ME) on PND 21. All mice were allowed to acclimate to the colony for one week (PND 21–27) prior to experimentation, where they were handled daily to allow for acclimation to experimenter manipulation. Animals were PND 28 of age at the beginning of the experiment. Animals were group-housed (4–5 per cage) with free access to food and water throughout the AIE exposure. Beginning at PND 47 (Experiment 1) or PND 68 (Experiment 2), mice were individually housed for intermittent two-bottle choice voluntary ethanol intake (described below). Food and water were available ad libitum throughout the experiment. All mice were maintained in a temperature and humidity-controlled room with lights on from 0700–1900 hr. Body weights were recorded daily during adolescence and weekly during adulthood. Animal care followed National Institutes of Health Guidelines under North Carolina Agricultural and Technical State University Institutional Animal Care and Use Committee approved protocols.

#### 2.2.1. Adolescent Intermittent Vapor Inhalation Chamber Exposure

Mice were exposed to repeated intermittent ethanol or air vapor for four two-day cycles from PND 28–42 for both experiments. Mice were exposed to ethanol vapor or air using protocols from previous work conducted in mice (Becker and Lopez, 2004; Carrara-Nascimento et al., 2013; Dhaher et al., 2008; Diaz-Granados and Graham, 2007; Griffin et al., 2009; Jury et al., 2017; Lopez and Becker, 2005; Maldonado-Devincci et al., 2014, 2016; Metten et al., 2010) and modified from Gass and colleagues (2014) for adolescent mice. On PND 28–29, 32–33, 36–37, 40–41, at approximately 1630 hr, mice were weighed and administered an intraperitoneal injection (0.02 ml/g) of the alcohol dehydrogenase inhibitor pyrazole (1 mmol/kg) combined with saline for control mice or combined with 1.6 g/kg ethanol (8% w/v) for ethanol-exposed mice and immediately placed in the inhalation chambers (23 in x 23 in x 13 in; Plas Labs, Lansing, MI). Mice remained in the vapor inhalation chambers for 16 hr overnight with room air (control group) or volatilized ethanol (ethanol group) delivered to the chambers at a rate of 10 liters/min. These procedures were designed to maintain blood ethanol concentrations (BECs) at 15\1–\275 mg/dl throughout the exposure period. Ethanol (95%) was volatilized by passing air through an air stone (gas diffuser) submerged in ethanol. The following morning (on PND 29, 33, 37, and 41), at 0900 hr, mice were removed from the vapor inhalation chamber and 50 μL of blood was collected from the submandibular space for BEC assessment. Mice were then returned to the home cage for 8 hr. On the evening of PND 29, 33, 37, and 41, mice were administered pyrazole and placed in the chamber overnight for 16 hours. On the morning of PND 30, 34, 38, and 42, mice were removed from the chambers and returned to the home cage. Mice remained undisturbed for two days between each cycle on PND 30-31, 34-35, 38-39. All blood samples were centrifuged at 5,000*g* and serum was collected and used to analyze blood ethanol concentrations using the AM1 blood alcohol analyzer (Analox Instruments, Lunenburg, MA).

#### 2.2.2. Abstinence

For Experiment 1, we left mice undisturbed from PND 42-46 in the home cage, except for regular cage maintenance. For Experiment 2, we left mice undisturbed from PND 42-67 in the home cage, except for regular cage maintenance. On PND 47 (Experiment 1) or PND 68 (Experiment 2), we transferred mice to individual cages to allow for acclimation to the intermittent two-bottle choice voluntary ethanol intake procedures.

#### 2.2.3. Voluntary Ethanol Intake

Beginning on PND 49 through PND 76 (Experiment 1) or PND 70 through PND 97 (Experiment 2), all mice were assessed for intermittent 24 hr voluntary ethanol intake using a two-bottle choice paradigm. Fresh tubes were presented to all mice daily. On PND 49 or PND 70, mice were introduced with two graduated sipper tubes containing water. Volumetric measurements were collected when tubes were introduced to the cages and on the following morning when tubes were removed. On PND 50 (Experiment 1) or PND 71 (Experiment 2), one water sipper tube was replaced with a tube containing 20% v/v unsweetened ethanol. Each two-day cycle was repeated for a total of 28 days, with intermittent presentation of the ethanol solution. Fresh tubes of water and/or ethanol were presented to all animals daily. Data were recorded daily for water and ethanol intake across the 28 days of intermittent ethanol access (PND 49-76-Experiment 1 or PND 70-97-Experiment 2).

The side of presentation of the ethanol and water sipper tubes were alternated to avoid the development of a side preference. Volumes were recorded to the nearest 0.1 ml. The difference in volume indicated the amount of ethanol consumed, and data are presented as grams of ethanol consumed per kilogram of body weight (g/kg) for the 24 hr session. Spillage was accounted for by placing the ethanol and water tubes in a similar holding cage unoccupied by a mouse. The difference calculated between the presentation and removal of the tube from the holding cage accounted for spillage and was subtracted from the daily difference calculated for each mouse. This procedure was repeated each day between 0900-1200 hr during the light cycle.

### 2.3 Open Field Test

Between PND 79-81 (Experiment 1) or PND 100-102 (Experiment 2), all mice were tested for a 60 min trial in the open field test. The animals were counterbalanced across groups for testing. Using a Plexiglas chamber (40.6 cm x 40.6 cm), mice were introduced to the open field testing arena (Kinder Scientific, Poway, CA) for 60 min to assess general exploratory behavior. Immediately upon removal from the arena, mice were returned to their home cage and returned to the colony. The open field arena was cleaned with 70% ethanol and allowed to dry completely before the next mouse was introduced. Data were quantified using Motor Monitoring Behavioral software (Kinder Scientific, Poway, CA).

### 2.4 Design and Analyses

Ethanol drinking and preference data were analyzed separately for each experiment for male and female mice using a two-factor mixed-model design ANOVA with Exposure (2; Air, Control) as a between subjects factors and Day as a repeated measure. Given the innate sex difference in ethanol intake in the control animals, data were normalized as a percent of control, with control animals set to zero and data for the ethanol-exposed mice expressed relative to zero. For the percent of control data, increased numbers indicate that ethanol-exposed mice consumed more ethanol than their air-exposed counterparts. The Percent of Control data for the ethanol-exposed mice were analyzed using a two-factor mixed model ANOVA with Sex (2; Male, Female) as a between subjects variable and Day as a repeated measure. All open field data were analyzed using a two-factor mixed model ANOVA with Exposure (2; Air, Control) and Sex (2; Male, Female) as between subjects factors. In the presence of significant interactions, Tukey’s and Sidak’s multiple comparisons post hoc tests were used where appropriate.

## 3. Results

### 3.1. Blood Ethanol Concentrations and Weight

All BEC data were analyzed using a three-way ANOVA for Sex by Cycle by Cohort given that all mice were exposed to the vapor inhalation chambers at the same age. There were differences between cohorts for blood ethanol concentrations as supported by a Cycle by Cohort interaction [F (3, 122) = 29.5, p < 0.0001] and a main effect of Cycle [F (3, 122) = 31.5, p < 0.0001]. BEC data are presented in Table 1 below. There were no sex differences between experiments. When comparing adolescent male ethanol-exposed mice between experiments, there were no differences in BECs at any given cycle. However, when comparing female ethanol-exposed mice between experiments, there were differences in BECs at each cycle.

**Table 1:** Blood ethanol concentrations (mg/dl) per cycle in male and female mice exposed to intermittent ethanol vapor inhalation. Data are presented as mean +/-SEM. Data are presented separately for each experiment. * indicates difference between male and female mice exposed to ethanol vapor inhalation.

| Experiment 1 |  |  | Experiment 2 |  |
| --- | --- | --- | --- | --- |
|  | Male | Female | Male | Female |
| **Cycle 1** | 278.9 ± 11.6 | 299.8 ± 11.6* | 238.5 ± 19.5 | 230.8 ± 15.3 |
| **Cycle 2** | 252.7 ± 28.9 | 225.7 ± 12.9* | 313.7 ± 22.7 | 319.0 ± 14.3 |
| **Cycle 3** | 214.1 ± 20.7 | 260.4 ± 11.9* | 165.6 ± 11.9 | 188.5 ± 8.6 |
| **Cycle 4** | 153.2 ± 22.9 | 159.3 ± 19.5* | 233.7 ± 19.2 | 272.0 ± 10.7 |

Specifically, during cycles 1 and 3, female mice exposed to ethanol during Experiment 1 showed higher BECs compared adolescent female mice exposed to ethanol vapor in Experiment 2.

During cycles 2 and 4, adolescent female mice in Experiment 2 showed higher BECs compared to adolescent female mice exposed to ethanol vapor during Experiment 1. Therefore, all subsequent analyses were conducted within each experiment.

#### 3.1.3 Experiment 1: Weight

When weight was analyzed as a three-factor mixed model ANOVA for Sex (2; Male, Female), Exposure (2; Air, Ethanol) and Day as a repeated measure, there was a significant Day by Sex interaction [F (23, 874) = 9.0, p < 0.0001] and Day by Exposure interaction [F (23, 874) = 3.2, p < 0.0001], and main effects of Sex [F (1, 38) = 81.71, p < 0.0001] and Day [F (23, 874) = 789.8, p < 0.001. Additionally, there was a trend for a main effect of Exposure [F (1, 38) = 3.2, p = 0.08]. Therefore, data were analyzed separately for each sex to assess ethanol-induced differences in weight across the experiment and are shown in Table 2. When male mice were compared for weight gain across the experiment, there were differences in weight between air-exposed and ethanol-exposed mice as supported by a Day by Exposure interaction [F (23, 414) = 2.7, p < 0.0001] and a main effect of Day [F (23, 414) = 416.4, p < 0.0001]. However, post hoc analyses failed to reveal any statistically significant differences at any given point between ethanol-exposed and air-exposed mice. There were no differences in weight between adolescent-ethanol-exposed and adolescent-air-exposed mice when all mice were voluntarily consuming ethanol using the two-bottle choice paradigm. When female mice were compared for weight gain across the experiment, there were no statistically significant differences in weight between air-exposed and ethanol-exposed mice, however there was a main effect of Day [F (23, 460) = 367.5, p < 0.0001].

**Table 2:** Weight (g) in male and female mice exposed to intermittent ethanol vapor inhalation during adolescence (PND 28-42) and two-bottle choice drinking and open field testing (PND 50-83). Data are presented as mean +/− SEM.

| **PND** | **Male**<br>**Air** | **Male**<br>**Ethanol** | **Female**<br>**Air** | **Female**<br>**Ethanol** |
| --- | --- | --- | --- | --- |
| **26** | 12.1 ± 0.2 | 11.4 ± 0.3 | 10.8 ± 0.3 | 10.5 ± 0.2 |
| **28** | 14.1 ± 0.3 | 13.8 ± 0.4 | 12.3 ± 0.3 | 12.1 ± 0.2 |
| **29** | 13.6 ± 0.4 | 12.7 ± 0.5 | 12.0 ± 0.3 | 11.5 ± 0.2 |
| **32** | 16.2 ± 0.3 | 14.7 ± 0.7 | 13.4 ± 0.3 | 12.8 ± 0.3 |
| **33** | 16.1 ± 0.3 | 14.1 ± 0.7 | 13.5 ± 0.3 | 12.8 ± 0.3 |
| **36** | 17.3 ± 0.4 | 16.0 ± 0.6 | 14.1 ± 0.2 | 13.8 ± 0.4 |
| **37** | 17.1 ± 0.4 | 15.4 ± 0.7 | 14.3 ± 0.2 | 13.4 ± 0.4 |
| **40** | 18.1 ± 0.4 | 16.8 ± 0.7 | 15.4 ± 0.3 | 14.5 ± 0.2 |
| **41** | 17.9 ± 0.4 | 16.3 ± 0.7 | 15.4 ± 0.2 | 14.3 ± 0.2 |
| **50** | 19.9 ± 0.5 | 19.5 ± 0.5 | 17.0 ± 0.2 | 16.7 ± 0.4 |
| **52** | 20.7 ± 0.5 | 20.1 ± 0.4 | 17.4 ± 0.2 | 17.4 ± 0.5 |
| **54** | 20.9 ± 0.4 | 20.6 ± 0.4 | 17.6 ± 0.2 | 17.5 ± 0.4 |
| **56** | 21.2 ± 0.4 | 20.9 ± 0.4 | 17.9 ± 0.1 | 17.9 ± 0.4 |
| **58** | 22.1 ± 0.6 | 21.2 ± 0.4 | 18.3 ± 0.2 | 18.2 ± 0.4 |
| **60** | 22.0 ± 0.4 | 21.7 ± 0.4 | 18.6 ± 0.2 | 18.3 ± 0.4 |
| **62** | 22.0 ± 0.4 | 21.8 ± 0.4 | 18.7 ± 0.2 | 18.6 ± 0.4 |
| **64** | 22.2 ± 0.5 | 21.9 ± 0.4 | 19.0 ± 0.2 | 18.5 ± 0.4 |
| **66** | 22.3 ± 0.4 | 22.1 ± 0.4 | 19.1 ± 0.2 | 18.7 ± 0.5 |
| **68** | 22.5 ± 0.4 | 22.4 ± 0.4 | 19.4 ± 0.2 | 18.8 ± 0.4 |
| **70** | 22.9 ± 0.4 | 22.7 ± 0.3 | 19.5 ± 0.2 | 19.1 ± 0.4 |
| **72** | 23.0 ± 0.4 | 22.8 ± 0.4 | 19.7 ± 0.2 | 19.3 ± 0.4 |
| **74** | 23.4 ± 0.4 | 23.2 ± 0.4 | 20 ± 0.3 | 19.7 ± 0.4 |
| **76** | 23.6 ± 0.5 | 23.5 ± 0.4 | 20.1 ± 0.2 | 19.8 ± 0.4 |
| **81-83** | 24.4 ± 0.4 | 23.1 ± 1.0 | 20.6 ± 0.2 | 19.9 ± 0.6 |

#### 3.1.4 Experiment 2: Weight

When weight was analyzed as a three-factor mixed model ANOVA for Sex (2; Male, Female), Exposure (2; Air, Ethanol) and Day as a repeated measure, there was a significant Sex by Exposure by Day interaction [F (15, 788) = 4.2, p < 0.0001], a Day by Exposure interaction [F (15, 788) = 3.4, p < 0.0001], Day by Sex interaction [F (15, 788) = 35.4, p < 0.0001], and main effects of Exposure [F (1, 54) = 4.3, p < 0.05], Sex [F (1, 54) = 162.4, p < 0.0001], and Day [F (15, 788) = 1604.0, p < 0.0001]. Therefore, data are shown in Table 3 and were analyzed separately for each sex to assess ethanol-induced differences in weight across the experiment.

**Table 3:**
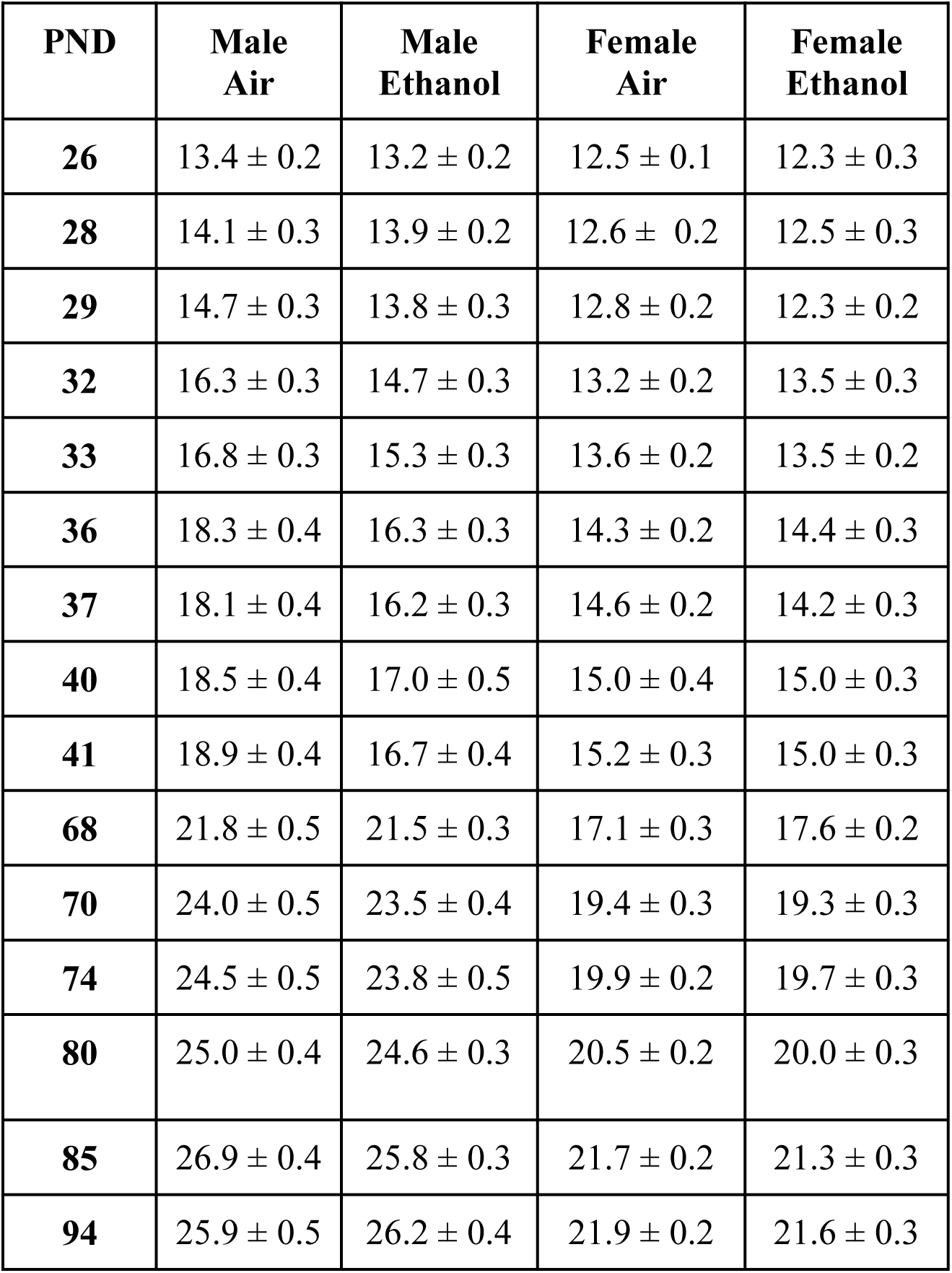
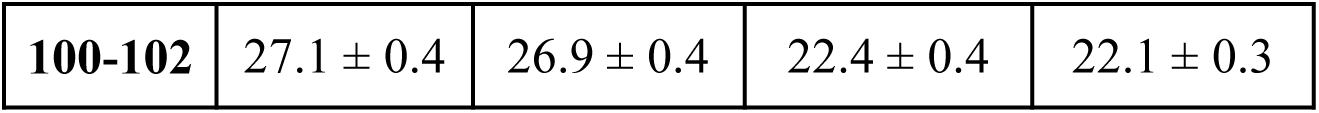
Weight (g) in male and female mice exposed to intermittent ethanol vapor inhalation during adolescence (PND 28-42) and two-bottle choice drinking and open field testing (PND 68-102). Data are presented as mean +/− SEM

When male mice were compared for weight gain across the experiment, there were differences in weight between air-exposed and ethanol-exposed mice on PND 32, 36, 37, and 41 (all p < 0.02) as supported by a Day by Exposure interaction [F (15, 387) = 5.0, p < 0.0001], a main effect of Exposure [F (1, 27) = 4.8, p < 0.05], and a main effect of Day [F (15, 387) = 757.4, p < 0.0001]. There were no differences in weight between male air-exposed and ethanol-exposed mice in adulthood when all mice were intermittently consuming ethanol. When air-exposed and ethanol-exposed female mice were compared for changes in weight, there were no differences between groups, but they did gain weight across the experiment as supported by a main effect of Day [F (15, 401) = 918.4, p < 0.0001].

### 3.2. Ethanol Consumption

#### 3.2.1. Experiment 1: Ethanol Consumption

Overall, there were changes in voluntary ethanol consumption during adolescence following one week since the last vapor exposure to ethanol. There were sex differences in ethanol consumption that varied as a function of exposure and day as supported by a Sex by Exposure interaction [F (1, 36) = 4.3, p < 0.05], and main effects of Sex F (1, 36) = 18.1, p < 0.0001] and Day [F (13, 468) = 6.7, p < 0.0001]. Given these sex differences in ethanol consumption, data were analyzed separately or male and female mice. As shown in Figure 1A adolescent ethanol-exposed male mice showed an overall increase in voluntary ethanol consumption compared adolescent air-exposed male mice (Figure 1A) as supported by a main effect of Exposure [F (1, 17) = 5.14, p < 0.05). Additionally, there was a main effect of Day [F (13, 221) = 2.9, p < 0.05). The Day by Exposure interaction failed to reach statistical significance. In adolescent female mice, AIE exposure did not alter voluntary ethanol consumption (Figure 1B). However, there was an increase in ethanol consumption among all groups across time as supported by a main effect of Day [F (13, 247 = 5.1. p < 0.001).

**Figure 1:**
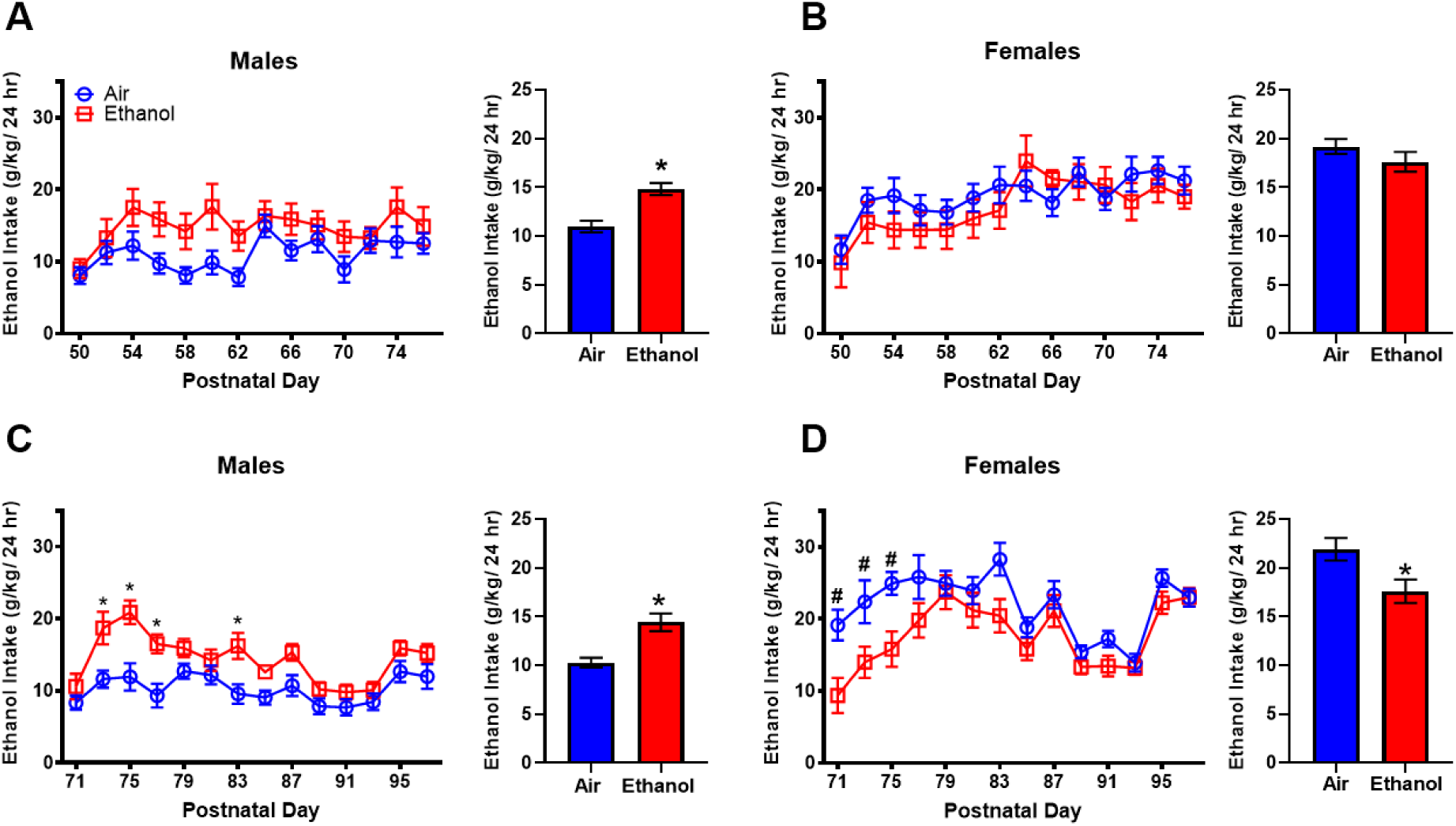
Intermittent voluntary ethanol consumption expressed as grams of ethanol consumed per kilogram of body weight (g/kg) across days (24 hr) following air or ethanol exposure during adolescence from PND 28-42. Panels A and B show ethanol consumption from experiment 1 in male and female mice from PND 50-74. Panels C and D show ethanol consumption from experiment 2 in male and female mice from PND 71-97. Data are presented as mean +/-SEM. * indicates difference between ethanol-exposed mice compared to air-exposed mice. # indicates female air-exposed mice drank more ethanol than female ethanol-exposed mice on that day.

#### 3.2.2. Experiment 2: Ethanol Consumption

Overall, there were long-term changes in ethanol consumption in adulthood following adolescent intermittent ethanol exposure in both male and female mice. There were sex differences in ethanol consumption that varied as a function of exposure and day as supported by a Day by Sex by Exposure interaction [F (13, 637) = 3.4, p < 0.0001], Sex by Exposure interaction [F (1, 49) = 17.4, p < 0.0001], Day by Sex interaction [F (13, 637) = 3.9, p < 0.0001], and main effects of Sex [F (1, 49) = 53.2, p < 0.0001] and Day [F (13, 637) = 19.9, p < 0.0001]. Given these sex differences in ethanol consumption, data were analyzed separately or male and female mice. As shown in Figure 1C, male mice that were exposed to ethanol during adolescence showed increased voluntary ethanol intake in adulthood on PND 73-77 (p < 0.005) and PND 83 (p < 0.01) compared to air-exposed controls, as supported by a significant Day by Exposure interaction [F(13, 312) = 2.9, p < 0.005] and a significant main effect of Exposure [F(1, 24) = 10.1, p < 0.005] and Day [F(13, 312) = 11.1, p < 0.0001]. In contrast, as shown in Figure 1D, female mice that were exposed to ethanol during adolescence showed less voluntary ethanol intake in adulthood on PND 71-75 (p < 0.05) compared to air-exposed controls, as supported by a significant Day by Exposure interaction [F(13, 325) = 2.0, p < 0.05] and a significant main effect of Exposure [F(1, 25) = 7.8, p < 0.05] and Day [F(13, 325) = 12.5, p < 0.0001].

Correlations between average BECS and average average ethanol intake (g/kg) were analyzed separately for each adolescent ethanol-exposed group. For adolescent ethanol-exposed male mice, there was a significant positive correlation (r = 0.74, p < 0.05) between average BEC and average g/kg ethanol consumption, but no significant correlation for adolescent ethanol-exposed female mice Experiment 1. There were no significant correlations average between average BECs and average g/kg ethanol consumption for either group in Experiment 2.

### 3.3. Preference Data

#### 3.3.1. Experiment 1

When ethanol preference data for experiment 1 (Figure 2A-B) were analyzed using a three-way ANOVA for Exposure (2; Air, Ethanol) and Sex (2; Male, Female) as between subjects factors and Day as a repeated measure, there were differences in overall preference for ethanol across days and between sexes, with a main effect of Day [F (13, 466) = 10.4, p < 0.0001] and Sex [F (1, 36) = 4.4, p < 0.05] and a trend for a Sex by Exposure interaction [F (1, 36) = 3.8 p = 0.06]. Therefore, data were analyzed separately for each sex using a two-factor ANOVA for Exposure and Day. As shown in Figure 2A for male mice, there was a main effect of Day [F (13, 221) = 3.9, p < 0.005], but no effect of Exposure. When daily water intake was averaged across the entire period of ethanol access, there was no difference in average daily water intake between adolescent air-exposed and adolescent ethanol-exposed male mice. Similarly, for female mice (Figure 2B), there was a main effect of Day [F (13, 247) = 8.2, p < 0.0001]. However, there were no differences in preference for ethanol between air-exposed and ethanol-exposed mice. There were also no differences in average daily water consumption between air-exposed and adolescent ethanol-exposed female mice.

**Figure 2:**
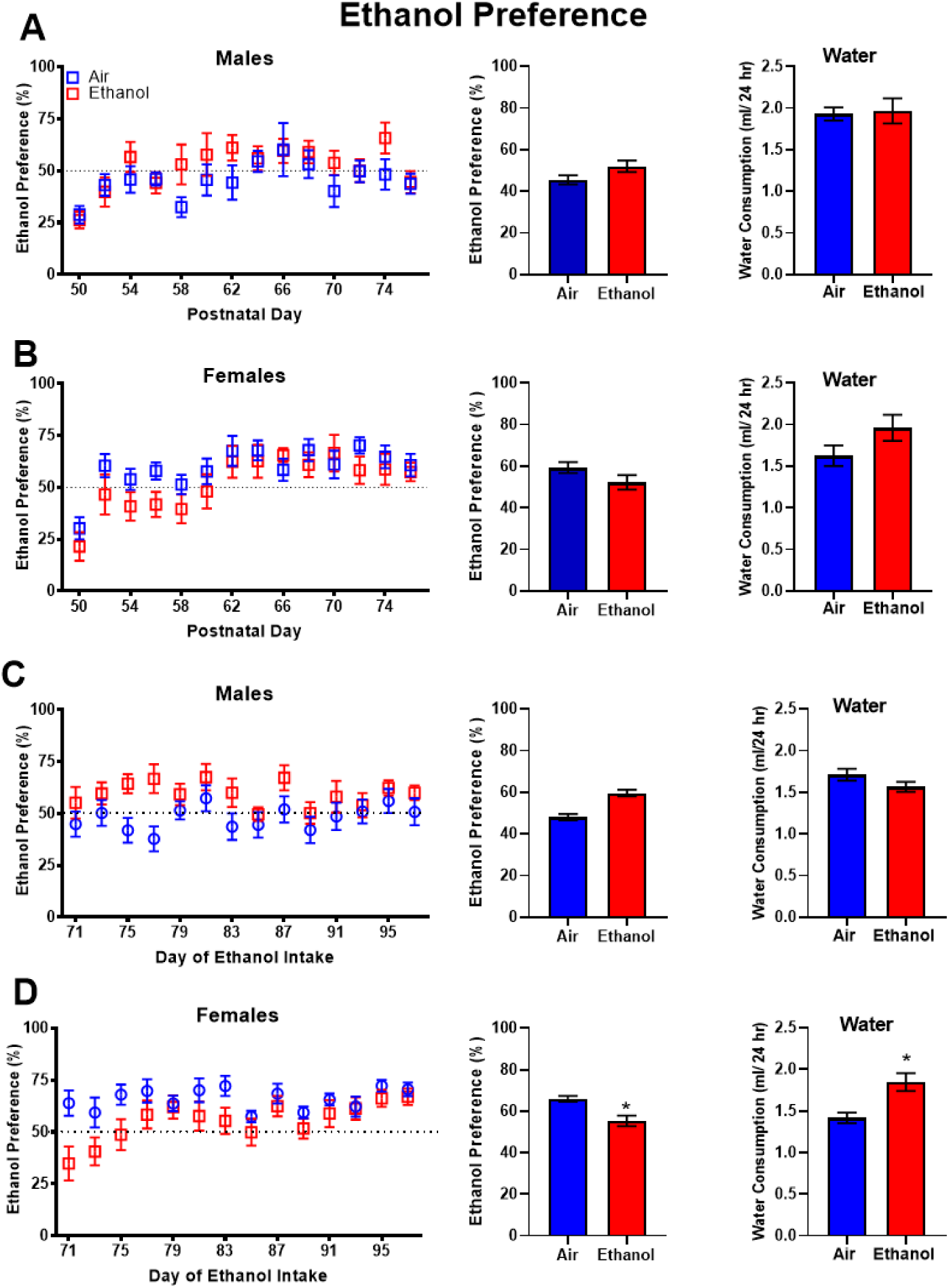
Preference for ethanol expressed as preference (%) computed as {(EtOH (ml)/ [EtOH (ml) + Water (ml))*100]} across days (24 hr) following air or ethanol exposure during adolescence from PND 28-42. Panels A and B show ethanol preference from experiment 1 in male and female mice from PND 50-74. Panels C and D show ethanol preference from experiment 2 in male and female mice from PND 71-97. Data are presented as mean +/-SEM. * indicates difference between ethanol-exposed mice compared to air-exposed mice.

#### 3.3.2. Experiment 2

When ethanol preference data for experiment 2 (Figure 2C-D) were analyzed using a three-way ANOVA for Exposure (2; Air, Ethanol) and Sex (2; Male, Female) as between subjects factors and Day as a repeated measure, there were differences in overall preference for ethanol across days and between sexes, with a main effect of Day [F (13, 637) = 4.9, p < 0.0001], a Sex by Exposure F (1, 49) = 8.4, p < 0.01] and a Sex by Day by Exposure interaction [F (13, 637) = 2.0 p < 0.05]. Therefore, data were analyzed separated for each sex using a two-factor ANOVA for Exposure and Day. For male mice, there was a trend for increased preference for ethanol in adulthood in the ethanol-exposed mice compared to control mice as shown in Figure 2C [Exposure (F (1, 24) = 4.1, p = 0.055)]. Additionally, there was a main effect of Day [F (13, 312) = 2.1, p < 0.05]. There were no differences in average daily water consumption between air-exposed and adolescent ethanol-exposed male mice. For female mice exposed to ethanol during adolescence, there was an overall decrease in preference for ethanol in adulthood (Figure 2D) as supported by a main effect of Exposure [F (1, 25) = 4.4, p < 0.05] and a main effect of Day [F (13, 325) = 4.3, p < 0.0001] and a trend for Day by Exposure interaction [F (13, 325) = 1.7, p = 0.06]. Average daily water intake was increased in adolescent ethanol-exposed female mice compared to adolescent air-exposed female mice.

### 3.4. Percent of Control Data

#### 3.4.1. Experiment 1

When sex differences in the control mice were examined, female control mice consumed approximately twice as much ethanol as compared to male control mice when data were expressed as g/kg as supported by a main effect of Sex [F (1, 19) = 26.4, p < 0.0001] and a main effect of Day [F (13, 247) = 4.7, p < 0.0005]. Given this innate sex difference in control mice, ethanol intake data cannot be directly compared to one another, therefore, ethanol intake data were transformed and expressed as a percent of the average respective sex air-exposed control. The ethanol intake data for the adolescent-air-exposed mice were averaged and set to 0 and individual daily ethanol intake data in the adolescent-ethanol-exposed mice were quantified relative to the averaged air-exposed control values [((adolescent ethanol-exposed / average adolescent air-exposed mean)*100)-100] to allow us to compare sex-differences in ethanol consumption for the adolescent-ethanol-exposed mice. Overall, adolescent ethanol-exposed male mice consumed more ethanol than adolescent ethanol-exposed female mice (Figure 3A) as supported by a Day by Sex interaction [F (13, 221) = 2.3, p < 0.01] and a main effect of Sex [F (1, 17) = 8.8, p < 0.01]. Specifically, adolescent ethanol-exposed male mice consumed between 73.3-91.3% more ethanol compared to ethanol-exposed female mice on PND 54-62 (all p < 0.03). On all other days, there were no differences in ethanol consumption between male and female mice.

**Figure 3:**
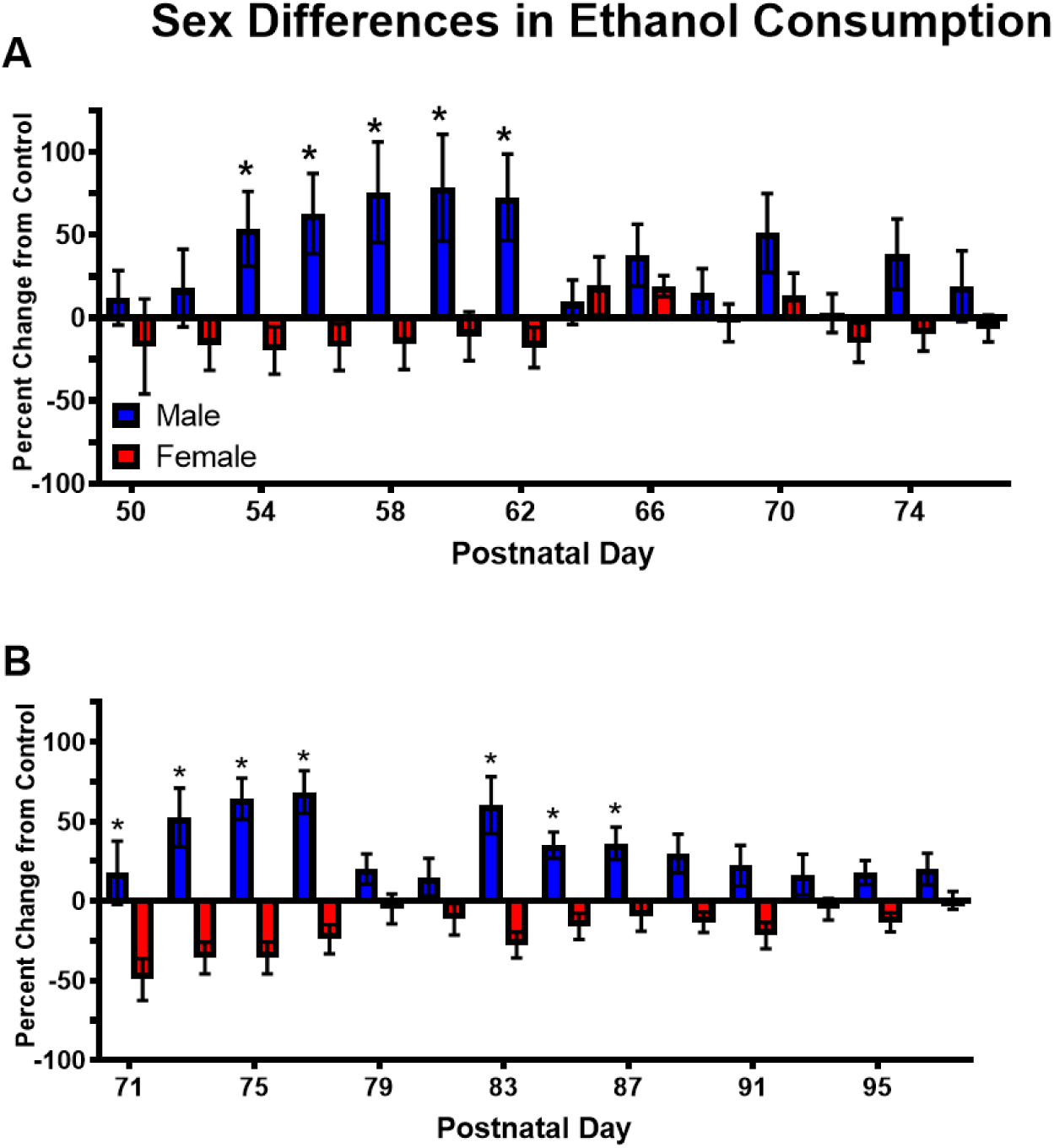
Voluntary ethanol consumption expressed as relative percent of control in ethanol-exposed mice across days (24 hr) following air or ethanol exposure during adolescence from PND 28-42. Control mice were set to 0 and ethanol intake was expressed relative to control, with higher scores indicating an increase in ethanol consumption relative to air-exposed mice and lower scores indicating a decrease in ethanol consumption relative to air-exposed mice. Panel A shows percent change from control ethanol consumption in ethanol-exposed mice from experiment 1 in male and female mice from PND 50-74. Panel B shows percent change from control ethanol consumption in ethanol-exposed mice from experiment 2 in male and female mice from PND 71-97. Data are presented as mean +/-SEM. * indicates males are significantly greater than females.

#### 3.4.2. Experiment 2

Sex differences in ethanol consumption were observed in control mice (main effect of Sex [F (1, 24) = 99.9, p < 0.0001]; main effect of Day [F (24, 312) = 7.8, p < 0.0001] and Day by Sex interaction [F (13, 312) = 2.6, p < 0.0002]. Therefore, as described above for Experiment 1, data were transformed and expressed as a percent of their respective sex control, with control mice set to 0 to allow us to compare sex-differences in ethanol consumption for the ethanol-exposed mice (Figure 3B). Overall, among the ethanol-exposed mice, males consumed more ethanol than females over the 28 days of ethanol consumption as supported by a significant main effect of Sex [F (1, 25) = 26.1, p < 0.0001] and Day [F (13, 325) = 2.2, p < 0.01], and a Sex by

Day interaction [F (13, 325) = 5.6, p < 0.0001]. Specifically, on PND 71-77, males consumed between 67.1-100% more ethanol than females (all p < 0.0005) and on PND 83-87, males consumed between 45.9-87.8% more ethanol than females (all p < 0.05). On all other days, there were no differences in ethanol consumption between males and females.

### 3.4. Open Field Test

#### 3.4.1. Experiment 1: Open Field

Data for the entire 60 min trial for the open first test were analyzed for total distance traveled and rearing. Overall, ethanol exposure decreased distance traveled compared to air-exposure during adolescence (Figure 4A) as supported by a main effect of Exposure [F (1, 38) = 4.4, p < 0.05] and a trend for a Sex by Exposure interaction [F (1, 38) = 3.7, p = 0.06]. There were no differences in total rearing frequency between any of the groups (Figure 4B).

**Figure 4:**
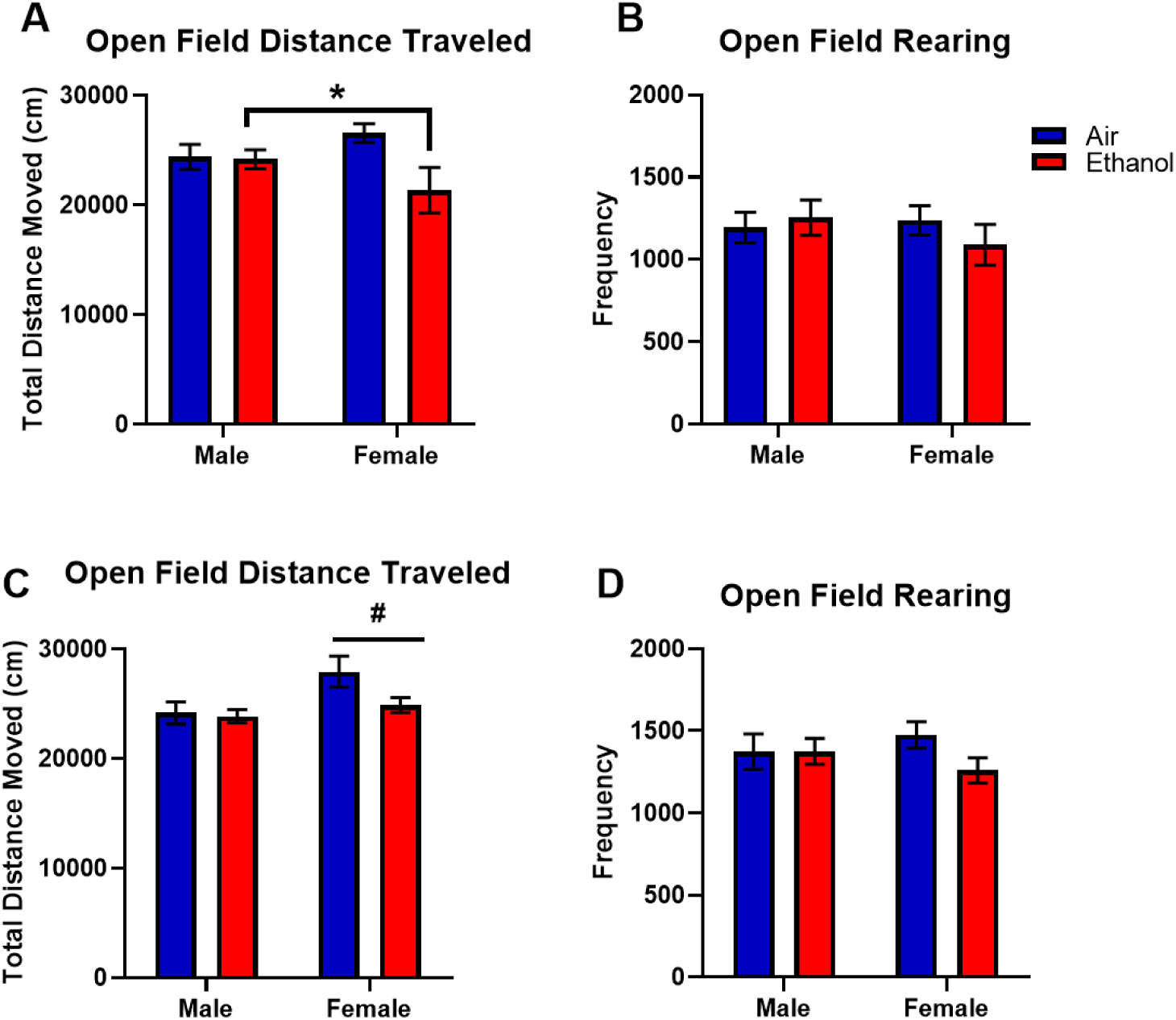
Behavior during the open field test for the entire arena following the 28 day voluntary ethanol drinking paradigm. Total distance traveled (cm) and rearing from experiment 1(Panels A and B) experiment 2 (Panels C and D) are shown. Data are presented as mean +/-SEM. * indicates ethanol-exposed mice are significantly greater than air-exposed mice. # indicates female mice significantly different from male mice.

#### 3.4.2. Experiment 2: Open Field

Overall, female mice showed greater distance traveled in the open field test compared to male mice (Figure 4C) as supported by a main effect of Sex [F (1, 49) = 6.02, p < 0.05].

Additionally, there was a trend for mice that were exposed to ethanol to show lower distance traveled compared to air-exposed control [Exposure (F (1, 49) = 2.96, p = 0.09)]. There were no changes in rearing behavior across the 60 minute trial (Figure 4D).

Correlations between average BECS and total TDM in the open field test were analyzed separately for each adolescent-ethanol-exposed group. There were no significant correlations between average BECs and total TDM in the open field test for either group in Experiment 1. For adolescent-ethanol-exposed male mice, there was a significant positive correlation (r = 0.70, p < 0.01) and for adolescent-ethanol-exposed female mice a significant negative correlation (r = 0.59, p < 0.05) between average BEC and total TDM in the open field test in Experiment 2.

When correlations for average g/kg ethanol consumption and total TDM in the open field test were conducted for all air-exposed and ethanol-exposed mice in Experiment 1, there were no significant correlations. When correlations for average g/kg ethanol consumption and total TDM in the open field test were conducted for all air-exposed and ethanol-exposed mice in Experiment 2, there was a significant positive correlation (r = 0.40, p < 0.05) in female, but not male, mice.

When correlations for average g/kg ethanol consumption and total rearing in the open field test were conducted for all air-exposed and ethanol-exposed mice in both experiments, there were no significant correlations.

#### 3.4.3. Experiment 1: Open Field Center Zone

As shown in Figure 5A, female mice that were exposed to ethanol during adolescence traveled less in the center zone compared to female mice that were exposed to air during adolescence as supported by a significant Sex by Exposure interaction [F (1, 38) = 4.0, p = 0.05]. There were no differences in time spent in the center zone between any of the groups (Figure 5B). Female mice exposed to ethanol during adolescence entered the center zone fewer times than those exposed to air during adolescence (Figure 5C) as supported by a significant Sex by Exposure interaction [F (1, 38) = 4.7, p < 0.05].

**Figure 5:**
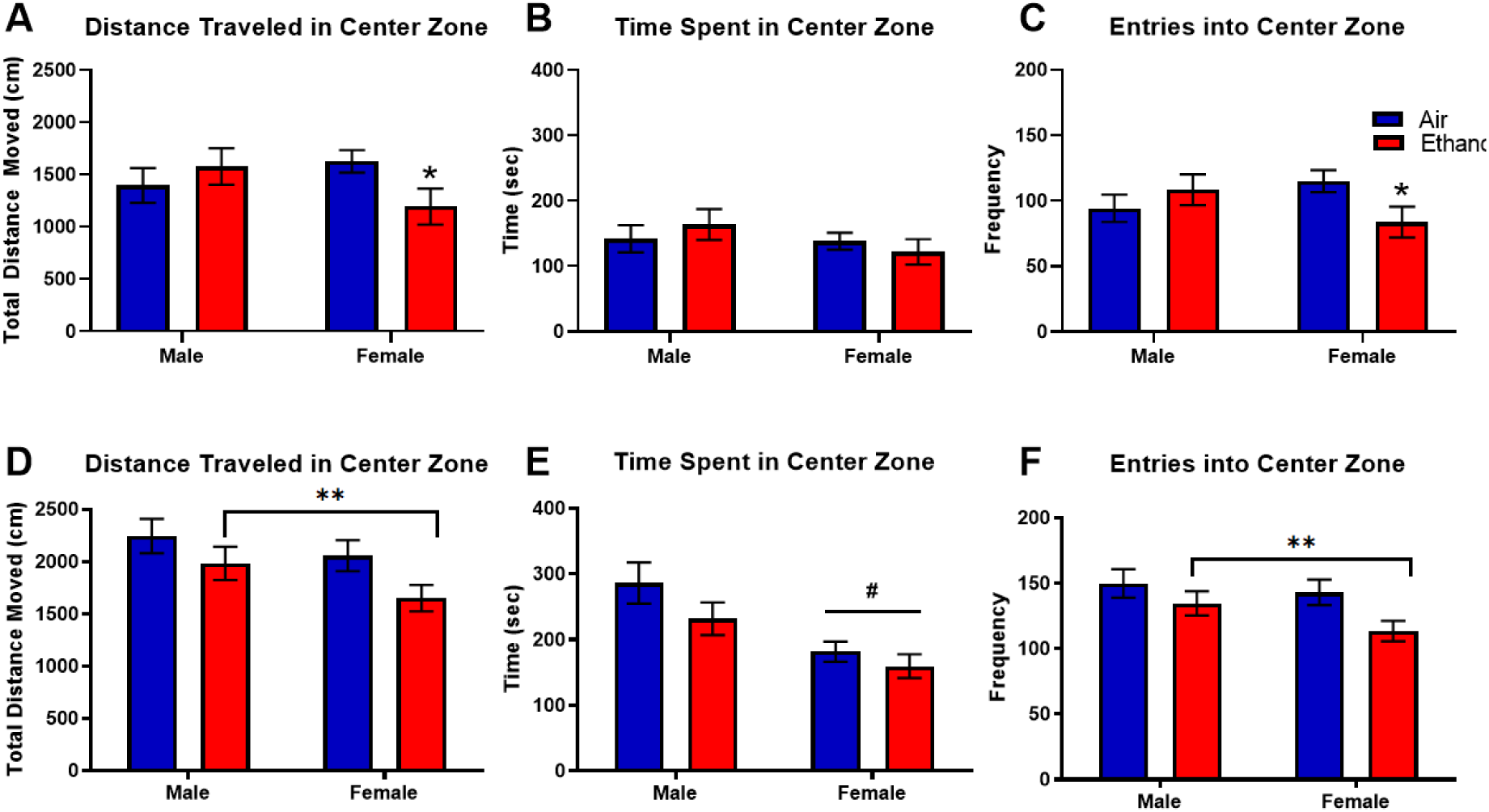
Behavior during the open field test for the center zone following the 28 day voluntary ethanol drinking paradigm between PND 79-81. Total distance traveled in centimeters for the center zone (Panel A), time spent (seconds) in the center zone (Panel B), and frequency of entries into the center zone (Panel C) for the entire 60 min trial. Data are presented as mean +/-SEM. * indicates female ethanol-exposed mice are significantly lower than female air-exposed mice. # indicates female mice significantly different from male mice. ** indicates ethanol-exposed mice are lower than air-exposed mice.

#### 3.4.4. Experiment 2: Open Field Center Zone

In general, AIE exposure decreased distance traveled and entries into the center zone, but did not change the amount of time spent in the center zone when mice were tested after a longer abstinence period. Specifically, mice exposed to ethanol during adolescence showed decreased movement in the center zone compared to mice exposed to air during adolescence (Figure 5D), as supported by a main effect of Exposure [F (1, 49) = 5.0, p < 0.05]. There was a trend for male mice to show greater distance traveled in the center zone compared to female miceg as supported by a trend for Sex [F (1, 49) = 3.0, p = 0.08]. As shown in Figure 5E, male mice spent more time in the center zone compared to female mice, regardless of exposure, as supported by a main effect of Sex [F (1, 49) = 14.6, p < 0.0005]. Figure 5F shows that AIE exposure decreased the number of entries into the center zone as supported by a main effect of Exposure [F (1, 49) = 5.64, p < 0.03].

When correlations for average g/kg ethanol consumption and center zone (1) TDM, (2) time spent, and (3) entries were conducted for all air-exposed and ethanol-exposed mice in Experiment 1, there were no significant correlations. When these same correlations were conducted for all air-exposed and ethanol-exposed mice in Experiment 2, there was a significant positive correlation in female mice for (1) TDM in the center zone and average ethanol consumption (r = 0.51, p < 0.01); (2) time spent in the center zone and average ethanol consumption (r = 0.47, p < 0.05); and (3) entries into the center zone and average ethanol consumption (r = 0.52, p 0.01). There were no significant correlations for male mice between any of these parameters in Experiment 2.

## 4. Discussion

The aim of this set of experiments was to determine if binge ethanol exposure during adolescence using intermittent vapor inhalation exposure would alter intermittent two-bottle choice ethanol consumption and open field behavior differently in male and female mice following one week of abstinence from ethanol exposure (during late adolescence) and in adulthood. We expected AIE vapor exposure to increase voluntary ethanol consumption compared to air-exposed mice both during late adolescence (Experiment 1) and during emerging adulthood (Experiment 2) in both male and female mice. Our data partially supported this hypothesis in that male mice that were exposed to AIE vapor during adolescence showed increased drinking in later adolescence and in young adulthood, an effect absent in female mice. AIE vapor exposure decreased center zone open field behavior in female mice during late adolescence and in young adulthood, an effect that was delayed until adulthood for male mice that were exposed to AIE.

### Ethanol Consumption

Our findings are consistent with recent work using chronic exposure in a drinking in the dark (DID) paradigm (Younis et al., 2019). Other work has shown that CIE exposure increased ethanol consumption in male but not female mice (Jury et al., 2017). Mid adolescence or between PND 44-51 has been identified as a transition period important for changes in ethanol consumption (Hargreaves et al., 2009), and this is the period during which animals in these experiments began the one week or four weeks withdrawal period. Consistent with this previous work that used intermittent access to ethanol (Hargreaves et al., 2009; Salguero et al., 2020; Tambour et al., 2008; Younis et al., 2019), in the present work we showed in both experiments that intermittent access to ethanol drinking increased ethanol intake during the initial part of the drinking paradigm, and this effect dissipated across the 28 days of ethanol consumption.

In female mice there was no effect of ethanol exposure on voluntary alcohol consumption during late adolescence, and we saw a transient decrease in voluntary ethanol consumption in adulthood. Others have shown that CIE exposure during adolescence did not change ethanol consumption in female mice, even when manipulating increasing concentrations of ethanol across the drinking access period (Jury et al., 2017). One proposed interpretation of these findings is that female mice may be protected from the effects of early ethanol exposure compared to male mice (Jury et al., 2017). It is clear we need more work to determine how ethanol exposure during adolescence changes later ethanol consumption in females.

Adolescent ethanol-exposed male mice did now show increased preference for ethanol during late adolescence in experiment 1 and showed a trend for increased preference for ethanol in young adulthood in experiment 2. This is similar to previous work where CIE-exposed male mice show an increase in ethanol consumption, but no change in preference for ethanol (Dhaher et al., 2008). For female mice that were exposed to ethanol during adolescence, there was no change in preference for ethanol during late adolescence (Experiment 1) and decreased preference for ethanol in emerging adulthood (Experiment 2), when compared to control air-exposed mice. Specifically, these mice showed an increase in average water consumption, which would account for the decrease in preference. These findings are similar to recent work done following CIE exposure in male and female adult mice and adolescent and adult female mice (Jury et al., 2017).

To account for the innate sex difference in ethanol consumption, where female control mice consumed more ethanol than male control mice, we transformed ethanol consumption data in the adolescent ethanol-exposed mice as a percent of control to allow us to compare relative differences in ethanol consumption between sexes. When data were expressed relative to control mice, we observed that adolescent ethanol-exposed male mice consumed more ethanol than adolescent ethanol-exposed female mice during late adolescence (experiment 1) and emerging adulthood (experiment 2,) primarily during the first half of the 28 day ethanol drinking paradigm and this effect dissipated across the second half of the ethanol drinking in both experiments.

### Open Field Behavior

During adolescence, there is lower sensitivity to alcohol induced motor impairments (Day et al., 2013), greater sensation seeking (Spear, 2016), and lower risk aversion (Casey and Jones, 2010). To assess potential consequences on the motor system from ethanol exposure during adolescence, we tested the mice in an open field test following 28 days of intermittent ethanol drinking. When mice from experiment 1 were tested in the open field test, the adolescent ethanol-exposed mice showed a modest decrease in overall distance traveled, an effect that appeared to be driven by the decrease in distance traveled in the adolescent ethanol-exposed female mice. When the mice were tested in later adulthood in experiment 2 following the 28 days of ethanol drinking, female mice showed greater distance traveled compared to male mice, regardless of ethanol exposure during adolescence. Previous research has shown that exposure to ethanol during adolescence and immediately testing rats for locomotor activation increased locomotor activity, but not when tested 30 minutes later (Acevedo et al., 2010). In the present work we tested mice in a drug-free state two days following the last access to ethanol drinking for 60 minutes. Future work should assess locomotor activation in response to an ethanol challenge administration.

The open field test is one of the most commonly used measures of anxiety-like behavior by using the center zone as a measure of thigmotaxis (Bolivar et al., 2000; Crawley and Bailey, 2008; Lipkind et al., 2004; Simon et al., 1994; Walz et al., 2016). Mice naturally have a phobia of bright open spaces, therefore a more anxious mouse would spend less time in the center zone of the open field. Using this test, more movement, entries, and time spent in the center zone are considered characteristics of less anxiety (Kulikov et al., 2008). We showed that the decrease in center zone ‘anxiety-like behavior’ emerged in female mice that underwent one week and four weeks withdrawal from adolescent ethanol exposure prior to intermittent ethanol drinking. The male mice only showed similar changes in behavior after four weeks prior to intermittent ethanol drinking. In experiment 1, adolescent ethanol-exposed female mice entered and traveled less distance in the center zone compared to similarly air-exposed female mice. In experiment 2, both male and female mice adolescent ethanol-exposed mice showed this behavioral profile in the center zone in the open field test. Regardless of exposure, female mice showed less time spent in the center zone compared to males. The reasons behind this sex difference are not clear. Using the center zone data as a crude measure for ‘anxiety-like’ behavior, we can interpret these findings as female mice exposed to ethanol during adolescence showing an ‘anxiety-like’ behavioral phenotype earlier compared to male mice.

Recently it was shown that binge drinking using the DID model increased anxiety-like behaviors in adult male but not female C57BL/6J mice following 36 hr withdrawal using other behavioral measures including marble burying, the light dark test, and the elevated plus maze (Rath et al., 2020). Other work using the CIE model has shown that ethanol exposure does not change anxiety-like behavior in C57BL/6J mice in the light/dark test (McCool and Chappell, 2015). Overall, our results in the present work examining ‘anxiety-like’ behavior from the open field test using center zone data show similar results to recent findings where female mice show a persistent anxiety-like phenotype following short-term and long-term withdrawal (Leonardo Jimenez Chavez et al., 2020). In future work, other behavioral paradigms to measure anxiety-like behavior should be utilized to gain better insight into this behavior following ethanol exposure during adolescence in both male and female mice.

In the present work, female adolescent ethanol-exposed mice showed decreased weight gain during the AIE exposure in experiment 1, an effect absent in male mice. However, in experiment 2, the opposite pattern was present where adolescent ethanol-exposed male showed deceased weight gain during the AIE exposure. Despite the differences in weight between air-exposed and ethanol-exposed mice during the adolescent exposure period, there were no differences in weight during the intermittent drinking period between groups in either experiment that would account for the observed differences. Variations in BECs could account for the differences in weight in the ethanol-exposed compared to the air-exposed mice. Repeated withdrawal from ethanol exposure may also be a stressor (Becker, 2017) and may have influenced weight gain during ethanol exposure. Cycles with higher BECs would likely induce more severe withdrawal from ethanol during AIE cycles. There were no differences in weight between exposure groups during the intermittent ethanol drinking when all mice were exposed to ethanol. The sex-dependent impact of repeated withdrawal on differences in weight should be explored in future work.

Blood ethanol concentrations were different between cycles and between experiments 1 and 2, but similar to previous work conducted using vapor inhalation in adolescent rats (Slawecki and Betancourt, 2002). The differences in BECs may have contributed to some behavioral differences observed between experiments. Other studies have also shown between cycle variations in BECs in several mouse strains across experiments (Lopez et al., 2017).

Although efforts are made to reduce variability in blood ethanol concentrations across experiments, there can still be differences between mice, between cohorts, and between cycles based on individual differences in mouse ethanol metabolism, the development of tolerance, and the functionality of equipment and environmental factors. The main differences in BECs between experiments were shown in female mice. Therefore, when making comparisons across cohorts, it is vital to equate BECs for female mice.

The present set of work was conducted in mice, whereas many previous studies using the AIE model used rats (Barker et al., 2017; Centanni et al., 2017; Ehlers et al., 2013; Gass et al., 2014; Nentwig et al., 2019; Slawecki et al., 2004; Trantham-Davidson et al., 2017; Walker et al., 2010). In those previous studies, pyrazole was not administered to the rats when assessing the impact of AIE on long-term changes in brain, ethanol consumption, and other behaviors. However, when using mice and vapor inhalation, pyrazole is regularly used (Becker and Lopez, 2004; Lopez and Becker, 2005; Maldonado-Devincci et al., 2016, 2014; Metten et al., 2010), which may account for some differences observed between rat and mouse studies. It is not clear how female mice respond to pyrazole in the vapor dependence model and may need to be addressed in future studies.

There have been conflicting results regarding binge ethanol exposure during adolescence and ethanol consumption in adulthood in rodent models. In the present set of experiments, male mice seem to be more sensitive to the effects of early ethanol exposure on voluntary alcohol consumption later in life. In previous work, when AIE vapor exposure occurred during adolescence, mice showed alterations in PFC-related decision making and behavioral control, hypoactivity in locomotion tests, increased anxiety-like behavior in the open field test, depressive-like behavior and lower body weight following exposure, and sex differences in negative affect (Centanni et al., 2017; Ehlers et al., 2013; Gass et al., 2014; Kasten et al., 2020; Nentwig et al., 2019; Slawecki et al., 2004; Walker et al., 2010). It is important to note that in the present work, the increased drinking behavioral phenotype in male mice began to emerge after the 7 day abstinence period during late adolescence and was more robust after the 27 day abstinence period in emerging adulthood. Together, these data show sex-dependent persistent changes in alcohol drinking, ‘anxiety-like’ behavior, and general exploratory behavior well into adulthood following AIE exposure. These findings suggest that male mice show persistent changes manifested as a change in ethanol-drinking phenotype based on their increased voluntary consumption in adulthood. Female mice appear to be more affected in locomotion and ‘anxiety-like’ behavior based on their reduced distance traveled and center-zone behavioral changes in the open field test. These data demonstrate a sexually divergent shift in ethanol consumption following binge ethanol exposure during adolescence and highlight sex-dependent vulnerability to developing substance use disorder in adulthood.

## Author Contributions

NIH, RCW, MJS, and AK collected the data. AMD, AMH, JGM, RCW, and NIH drafted the manuscript. AMD designed the experiments and analyzed the data. AMD, RCW, JGM, AMH, and NIH edited the manuscript.

## Other Acknowledgements

The authors would like to thank Myracle C. Jones, Ashley M. Davis, Victoria Robinson, Destiny M. Belton, and Kristoni Barnes for their expert technical assistance.

